# Ecotone formation through ecological niche construction: the role of biodiversity and species interactions

**DOI:** 10.1101/740282

**Authors:** Kevin Liautaud, Matthieu Barbier, Michel Loreau

## Abstract

Rapid changes in species composition, also known as ecotones, can result from various causes including rapid changes in environmental conditions, or physiological thresholds. The possibility that ecotones arise from ecological niche construction by ecosystem engineers has received little attention. In this study, we investigate how the diversity of ecosystem engineers, and their interactions, can give rise to ecotones. We build a spatially explicit dynamical model that couples a multispecies community and its abiotic environment. We use numerical simulations and analytical techniques to determine the biotic and abiotic conditions under which ecotone emergence is expected to occur, and the role of biodiversity therein. We show that the diversity of ecosystem engineers can lead to indirect interactions through the modification of their shared environment. These interactions, which can be either competitive or mutualistic, can lead to the emergence of discrete communities in space, separated by sharp ecotones where a high species turnover is observed. Considering biodiversity is thus critical when studying the influence of species-environment interactions on the emergence of ecotones. This is especially true for the wide range of species that have small to moderate effects on their environment. Our work highlights new mechanisms by which biodiversity loss could cause significant changes in spatial community patterns in changing environments.

## 1 Introduction

Whether species composition changes gradually, or forms discrete zones along environmental gradients has been the subject of a long-standing debate in ecology (Clements, 1916; Gleason, 1926; Braun-Blanquet, 1928; Hedberg, 1955; McIntosh, 1967). Observational studies have found both gradual (Whittaker, 1956; Vazquez G. and Givnish, 1998; Ellison et al., 2010; Lieberman et al., 1996) and discrete patterns (Kitayama, 1992; Hemp, 2006; Tuomisto and Ruokolainen, 1994; Kessler, 2000). Rapid changes in community composition along gradients, also termed ecotones (Kent et al., 1997), have been observed in a wide range of ecosystems, such as alpine treelines (Germino et al., 2002), tropical mountain forests (Martin et al., 2007) and coastal environments (Sternberg et al., 2007; Walker et al., 2003). Hereafter, a transition will be termed “rapid” when its scale is much smaller than the spatial scale of the landscape, even though the transitional area may show mixing of species.

While rapid changes can be blurred by species dispersal (Liautaud et al., 2019) or stochasticity in nature, it is important to understand the theoretical conditions under which rapid community changes can emerge. These rapid changes in species composition can coincide with rapid changes in environmental conditions, such as the frost line (Kitayama and Mueller-Dombois, 1992) or discontinuities in edaphic conditions (Tuomisto and Ruokolainen, 1994; Kessler, 2000). In these cases, it is often assumed that changes in abiotic conditions are responsible for the change in species composition (McIntosh, 1967; Kent et al., 1997). This assumption is supported in many cases, but it may obscure the possibility that, in other settings, the two boundaries emerge together from the influence of species on their abiotic environment. The mechanisms that can lead to such transitions are poorly known, and in particular the respective contributions of species-environment feedbacks and interspecific interactions.

Species that are able to modify their abiotic environment are often called “ecosystem engineers” (Jones et al., 2010). Classical examples range from beavers that impact water flow and habitat heterogeneity (Wright et al., 2002), to cushion alpine plants that buffer extreme temperatures and increase soil moisture (Badano et al., 2006). Ecological niche construction is a particular case in which engineers modify the environment to their own benefits (Kylafis and Loreau, 2008, 2011), creating a feedback with the environment (an example in which engineers can instead create succession is presented in the Appendix). This ecological process should be distinguished from the related concept of niche construction in evolutionary theory in which we would also expect species traits to evolve over time (Odling-Smee et al., 2003, 1996). Examples of ecological niche construction range from plant-water feedbacks in arid environment (Dekker et al., 2007) to increases in nutrient inputs by trees in tropical ecosystems (De longe et al., 2008). Such feedbacks can govern species distributions (Wilson and Agnew, 1992), particularly under harsh environmental conditions (Kéfi et al., 2007; Gilad et al., 2004; Meron et al., 2004; von Hardenberg et al., 2001), and lead to the emergence of ecotones (Bearup and Blasius, 2017; Jiang and DeAngelis, 2013). Classical studies on ecosystem engineers, however, have generally focused on the effects of a particular species having strong effects on the abiotic environment (Jones et al., 2010; Bouma et al., 2010; Prugh and Brashares, 2012). But many more species have small or moderate impacts on their environment. Such species, which are often neglected individually, might substantially affect their environment when aggregated. Furthermore, previous studies have scarcely explored what types of interactions can arise between multiple species that engineer their shared environment. We thus propose to focus on the role of diversity and species interactions in the emergence of ecotones through ecological niche construction.

Biodiversity can have two main effects on the emergence of species-environment feedbacks: a cumulative effect of species number, and a heterogeneity effect due to variations in species’ preferences and engineering ability. Cumulative effects are similar to complementarity in biodiversity-ecosystem functioning relationships (Loreau and Hector, 2001; Hooper et al.). The fact that species coexist with weak or no competition implies the existence of different niches, i.e. other factors beyond the environmental preference modelled here. This cumulative effect arises when there is no single identifiable engineer, but where community acts collectively to create an ecotone. A potential example is the occurrence of ecotones between mangroves and hardwood forests, where several mangrove tree species can modify water salinity in synergy (Sternberg et al., 2007). In contrast, the heterogeneity effect of biodiversity arises when there are differences in species’ preferred environmental states. We investigate the effect of these differences on emergent competition or facilitation between ecosystem engineers, and how this could play a role in ecotone emergence.

In this study, we build a theoretical model that couples the dynamics of a community and of its abiotic environment to assess the role of ecosystem engineers and of their diversity in the emergence of ecotones in space. In our model, ecotones are represented by abrupt changes, including discontinuities. In the presence of multiple interacting species, we show that ecological niche construction can lead to the emergence of indirect interspecific interactions -which can be either positive or negative - through environmental modifications. Similarly, we show that even species with different preferences can act synergistically as a single community. We then assess the consequences of these different interaction types for community patterns in space, and identify the conditions under which ecotone formation is predicted to occur.

## 2 Model and methods

### 2.1 >Species growth and niche construction

We model the dynamics of a community of *n* species, each of which obeys a logistic growth along a gradient of an arbitrary environmental factor *E*. We consider independent locations along this environmental gradient, assuming no fluxes between the locations ^1^. For a given location *k*, the population dynamics of species *i* is given by:

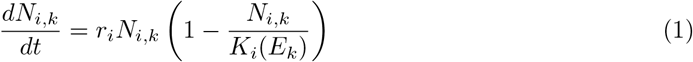

where *E*_*k*_ represents the value of the environmental factor at location *k*, 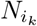 is the abundance of species *i* at that location, and *r*_*i*_ is its intrinsic growth rate, assumed to be equal for all species, *r*_*i*_ = *r*. The fundamental niche of each species is defined by its carrying capacity *K*_*i*_(*E*), which is assumed to depend on the environmental value *E* according to a Gaussian function:

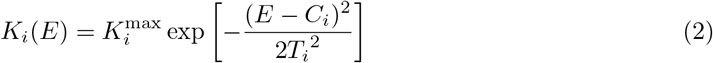

The classical Hutchinsonian niche (Hutchinson, 1957) would instead be defined in terms of growth rate, but these two assumptions are equivalent in the case of logistic growth as considered here. The above function is characterized by the species’ fundamental niche centre *C*_*i*_, i.e. the value of the environmental factor for which its carrying capacity reaches its maximum value 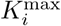, and its tolerance range *T*_*i*_. This unimodal, continuous distribution ensures a gradual response of each species to changes in the environment.

At each location *k* on the gradient, the environmental factor has a distinct physical baseline value *B*_*k*_ representing its state in the absence of environment modification. Species, however, can affect the environmental value *E*_*k*_ by pushing it toward their preferred value *C*_*i*_ at a maximum rate *m*_*i*_, which we call the niche construction rate. These species will be called “ecosystem engineers”. The environment tends to return spontaneously to its baseline value *B*_*k*_ at a rate *μ*. The dynamics of the environmental factor at location *k* is therefore:

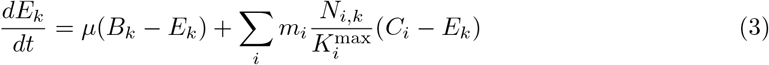

where abundance *N_i,k_* is rescaled by its maximum 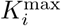 so that *m*_*i*_ is the maximum rate at which species *i* can affect the environment. In this study, we assume that species’ carrying capacities are only influenced by a single factor *E*, although we recognize that many abiotic factors can also affect *K* in nature. The presence of direct competition between species can also have an influence on species distributions in space (Liautaud et al., 2019), we describe this case in Appendix A3. In this simplified model, the only role played by growth rates is to determine how fast species reach their carrying capacities, and which equilibrium is reached from given initial conditions when there are multiple equilibria. The identification of alternative equilibria in described in the next section.

### 2.2 Potential landscape and alternative equilibria

To predict the long-term spatial patterns created by dynamics (1) and (3), we propose a simple method for finding their equilibria at each location *k* along the gradient. This method is based on the notion of potential landscape, whose role in ecology was pioneered by Holling (1973).

Let us consider a local community at a given location *k* with baseline environmental state *B*_*k*_. If species population dynamics are much faster than that of the environment (*r* ≫ max(*m_i_, μ*)) we expect that species quickly reach their carrying capacity for a given environment value, *N_i,k_* = *K*_*i*_(*E*_*k*_), while *E*_*k*_ changes over longer time scales according to:

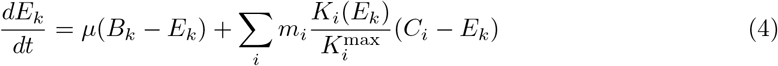

We show in the Appendix A2.1 that this can be expressed as a gradient descent dynamics,

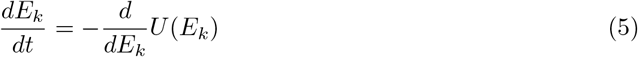

where *U* (*E*_*k*_) is a potential function. This equation imposes that, from any initial condition, the variable *E*_*k*_(*t*) always moves over time toward the closest minimum of *U* (*E*), and then stays there at equilibrium. This potential takes the form:

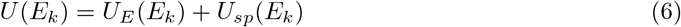

where *U*_*E*_(*E*_*k*_) represents the contribution of abiotic processes returning the environment to its baseline state, with

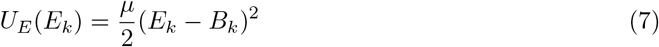

and *U*_*sp*_(*E*_*k*_) represents the species’ contribution

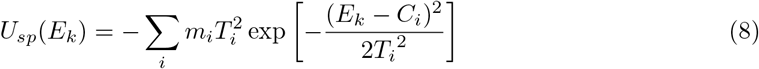

which we illustrate in Fig.1 for a single species. The relative effect of abiotic and biotic factors in encapsulated in the ratio:

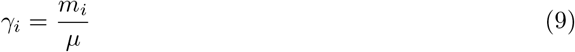

such that niche construction is weak for *γ* ≪ 1 and strong for *γ* ≫ 1. This parameter will be termed “niche construction strength”.

This potential landscape provides an intuitive interpretation of the action of engineer species. In the absence of niche construction (*m*_*i*_ = 0), the only minimum of *U* (*E*_*k*_) is at the physical baseline *E*_*k*_ = *B*_*k*_. When present, ecosystem engineers “dig” in that landscape, creating wells of width *T*_*i*_ centered on their preferred value *C*_*i*_. As we see in Fig.1, weak engineering only slightly displaces the equilibrium, while strong engineering can create an alternative equilibrium, or even overcome abiotic dynamics entirely.

**Figure 1:**
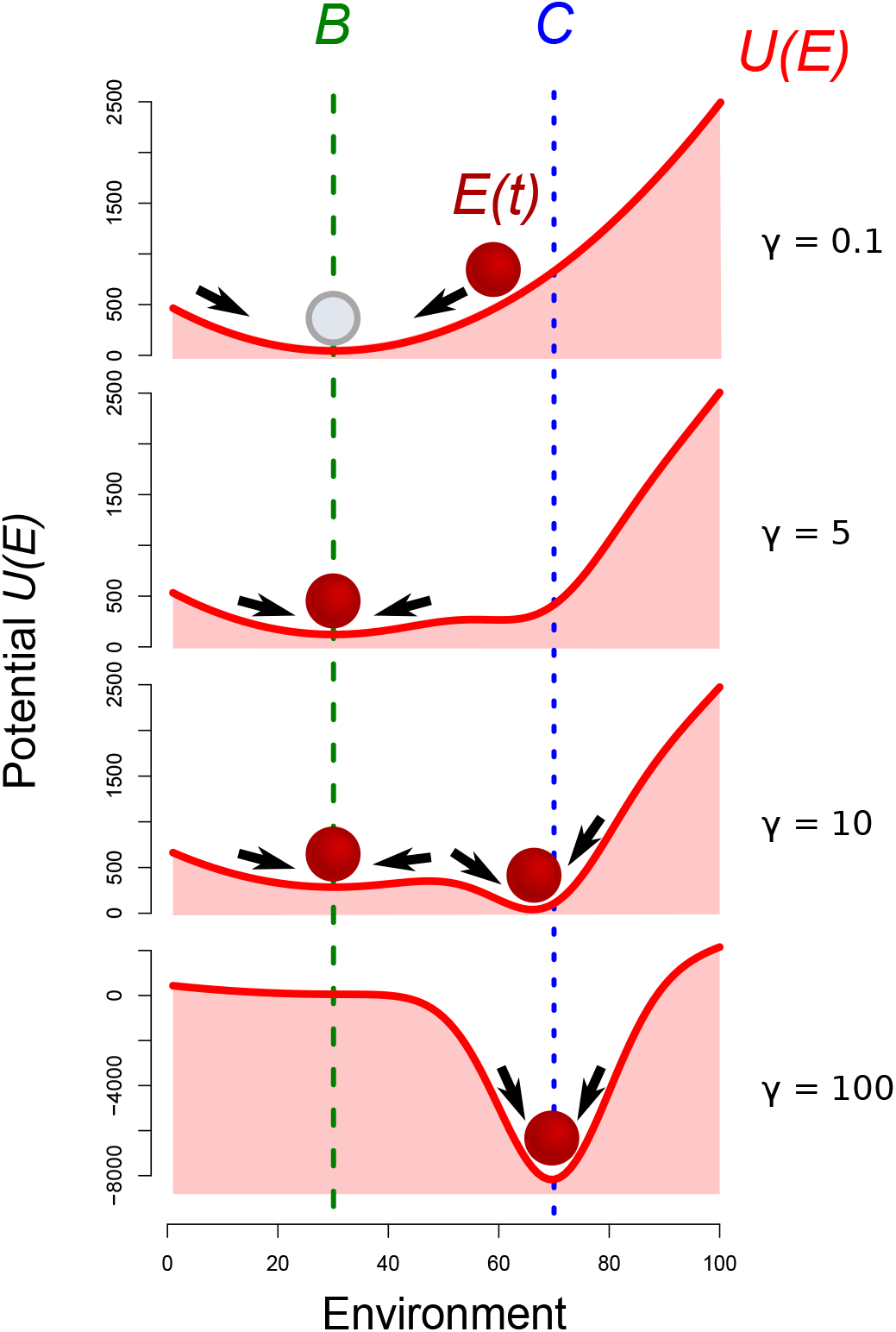
Representation of the environment as a potential under the action of physical processes and an ecosystem engineer. The ball representing the environmental state *E*(*t*) follows the arrows down the slope until it reaches an equilibrium value, corresponding to a minimum of the potential function *U* (*E*) (denoted by the solid curve). *B* is the baseline environment value, and *C* is the species’ environmental optimum. Four parameter conditions are depicted, from weak engineering (*γ* = 0.1) to strong engineering (*γ* = 100).

We also show in the Appendix A2.1 that, for arbitrary values of the rates *r*, *m*_*i*_ and *μ*, the dynamics of *E*_*k*_(*t*) become more complex than a gradient descent (i.e. the function *U* (*E*_*k*_) can increase for part of the time), but all possible equilibria are still given by the minima of the potential *U* (*E*_*k*_) defined in (6).

### 2.3 Numerical simulations

In the presence of a single ecosystem engineer, the niche construction strength (*γ*) is expected to be the main driver of the dynamics. We thus study the influence of this parameter on the shape of potential landscape, and the consequences for species’ distribution in space.

In diverse communities, the similarity of species in their ressource use or environmental requirements has been shown to influence species interactions (Abrams, 1983; MacArthur and Levins, 1967; Levin, 1970), and species distribution in space (MacArthur, 1972). Therefore, we study how the difference in the environment optimum of the various species (Δ*C*) and the niche construction strength (*γ*), can influence the nature and intensity of species interactions (*I*) in a two-species system. To do this, we compute the abundance of a species 1 when alone (*N*_1*a*_), or in the presence of a second species 2 (*N*_1*b*_), for different values of (*γ*, Δ*C*). We use the relative change in the abundance of species 1 as a measure of the net effect of species 2 on species 1:

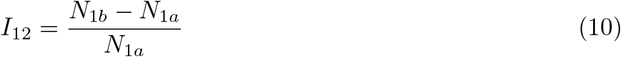

In our study, the two species have equal niche construction abilities, but distinct environment optima. In the case where bistability is observed, we only study the equilibrium for which species 1 predominates (*C*_1_ = 40, *C*_2_ = *C*_1_ + Δ*C*, *E*_*t*=0_ = *B* = 50). We then extend these results to a larger number (*S*) of engineer species.

To address the role of these different factors - (*γ*,Δ*C*, *S*) - on community pattern in diverse communities, we study an environmental gradient of 101 cells ranging from *k* = 100 to *k* = 200 in arbitrary units, with a step size of 1. The baseline value of the environment gradually increases along the gradient, as *B*_*k*_ = *k*. The centres of the fundamental niches of the various species, *C*_*i*_, are randomly assigned following a uniform distribution between 0 and 300, so that species may have their niche centre in or outside the studied zone initially.

The model is run independently on each cell. The initial value of the environment at each location equals its baseline value(*E*_*k*_(*t* = 0) = *B*_*k*_). For all simulation results in the main text, species were given equal maximal carrying capacity *K*^max^ = 1 and tolerance range *T ≤* 10. Environmental return rate is set to *μ* = 1, and species intrinsic growth rate is set to *r* = 10. Under these conditions, with *r ≫ μ*, species quickly reach their carrying capacity, with *N_i,k_* = *K_i,k_*(*E*) (see 2.2). Initial species abundances are set equal for all species in all locations. We run the model with different values of the different parameters of interest (*γ*, Δ*C*, *S*) until *t* = 1000, and verify that the equilibrium is reached.

## 3 Results

### 3.1 Effects of niche construction strength on local equilibria

In the case where niche construction is weak (*γ* = 0.1, Fig. 1), the dynamics goes towards the environmental baseline value *B*. However, when the niche construction strength of a species increases (*γ* = 5), it becomes able to influence the environment. With increasing niche construction, the species becomes able to create an alternative stable equilibrium, which corresponds to an environment value close to its optimum (*γ* = 10). For a very high niche construction ability (*γ* = 100), the species environment optimum becomes the single stable equilibrium in the system.

### 3.2 Engineer similarity, attractors and species interactions

Here we study the influence of the difference in engineers’ environment optima (Δ*C*) on the potential landscape. For 2 species with a high niche construction rate (*γ →* +*∞*, Fig. 2, a) Δ*C* determines the number of attractors in the system. We can calculate a threshold *θ* of Δ*C* that separates cases in which species’ contributions to the potential (*U*_*sp*_, Eq 6) create a single attractor, from cases where two attractors are observed. When Δ*C > θ*, there are two minima in *U*_*sp*_. As we have assumed that the abiotic contribution *U*_*E*_(*E*) is negligible, the species create distinct minima in the potential *U* (*E*) (red curve) that correspond to distinct attractors (i.e alternative stable states), in which the environment is optimal for either of the two species (Fig. 2, a, I). By contrast, when Δ*C < θ*, there is a single minimum in *U*_*sp*_. In this case, the two species create a common well in the potential landscape, which corresponds to a single equilibrium in between the two species’ optima (Fig. 2, a, II). We show in the Appendix A2.2 that *θ* = 2*T* for species with equal tolerance ranges *T* and maximal carrying capacities *K*_*max*_.

**Figure 2:**
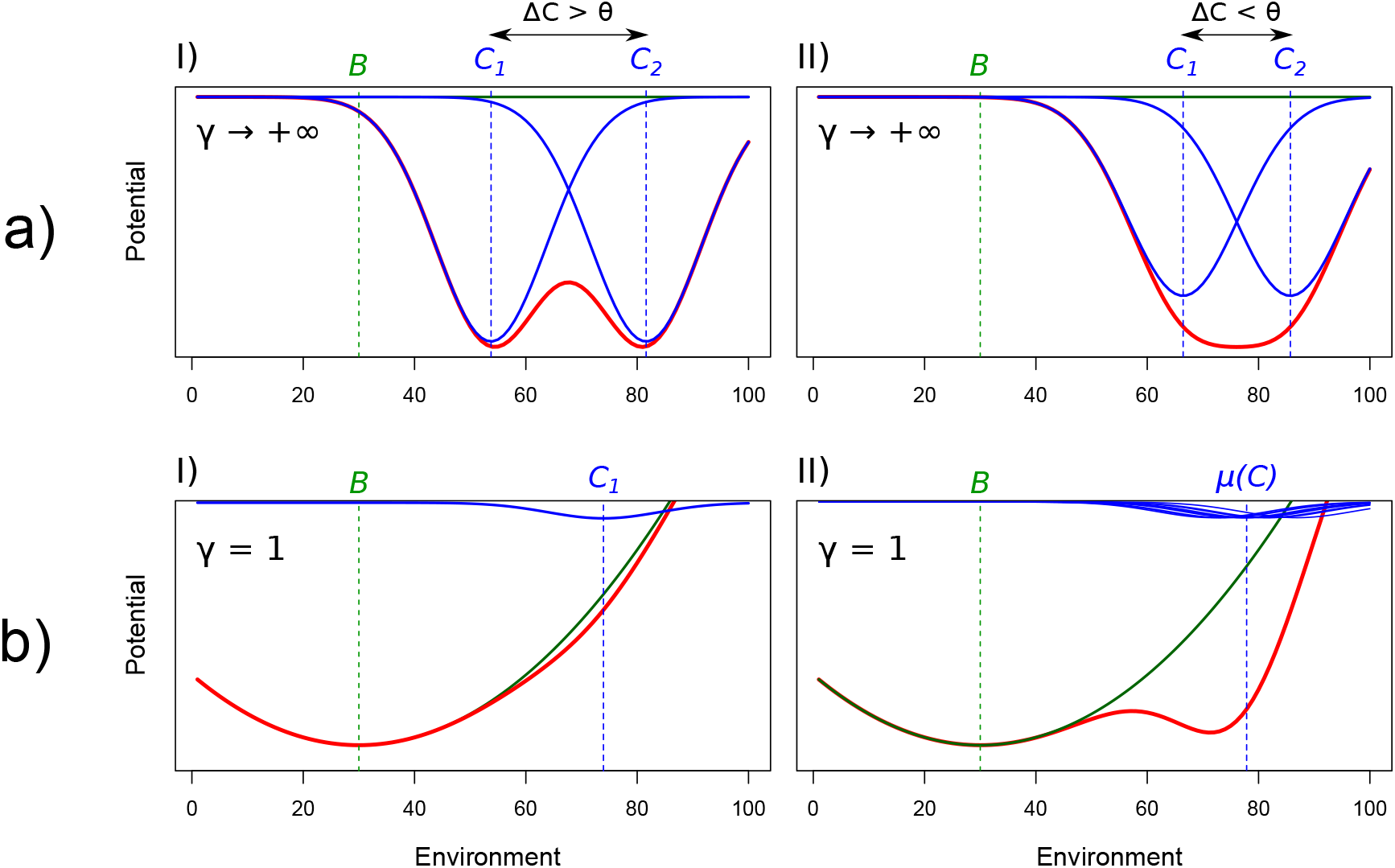
Influence of the similarity in species’ environmental optima (Δ*C*) and diversity (*S*) on the potential landscape. Blue and green curves show the contributions of species (*U*_*sp*_(*E*)) and environment (*U*_*E*_(*E*)), respectively, to the potential *U* (*E*) (red curve). a): Influence of strong ecosystem engineers (*γ* = +∞) on the potential landscape for two values of optimum similarity Δ*C*. *θ* represents the threshold in Δ*C* that separates cases in which species’ contribution to the potential (*U*_*sp*_(*E*)) show one or two minima. b): Influence of diversity in engineering species on the potential landscape for two levels of diversity: *S* = 1 (I) and *S* = 10 (II), for low niche construction strength (*γ* = 1).

The similarity (Δ*C*) of engineers therefore influences the nature and intensity of species net interactions. When niche construction is weak and the similarity in environmental optima is high, the abundance of species 1 is increased when associated with species 2 (Fig. 3, red). The relative increase in species 1’s abundance in association with species 2 can reach 8% when compared with its abundance when alone, indicating a positive net interaction between the two species (*I* > 0). By contrast, when niche construction is high and dissimilarity in environment optima is high, species 1 has a lower abundance in the presence of species 2 (Fig. 3, blue, indicating a negative net interaction (*I* < 0)). The relative decrease in the abundance of species 1 in the presence of species 2 can reach more than 30%, and is maximal for Δ*C ≈ θ*. For a given niche construction rate *γ*, indirect interactions can thus be alternatively positive or negative, depending on the species’ similarity Δ*C*.

**Figure 3:**
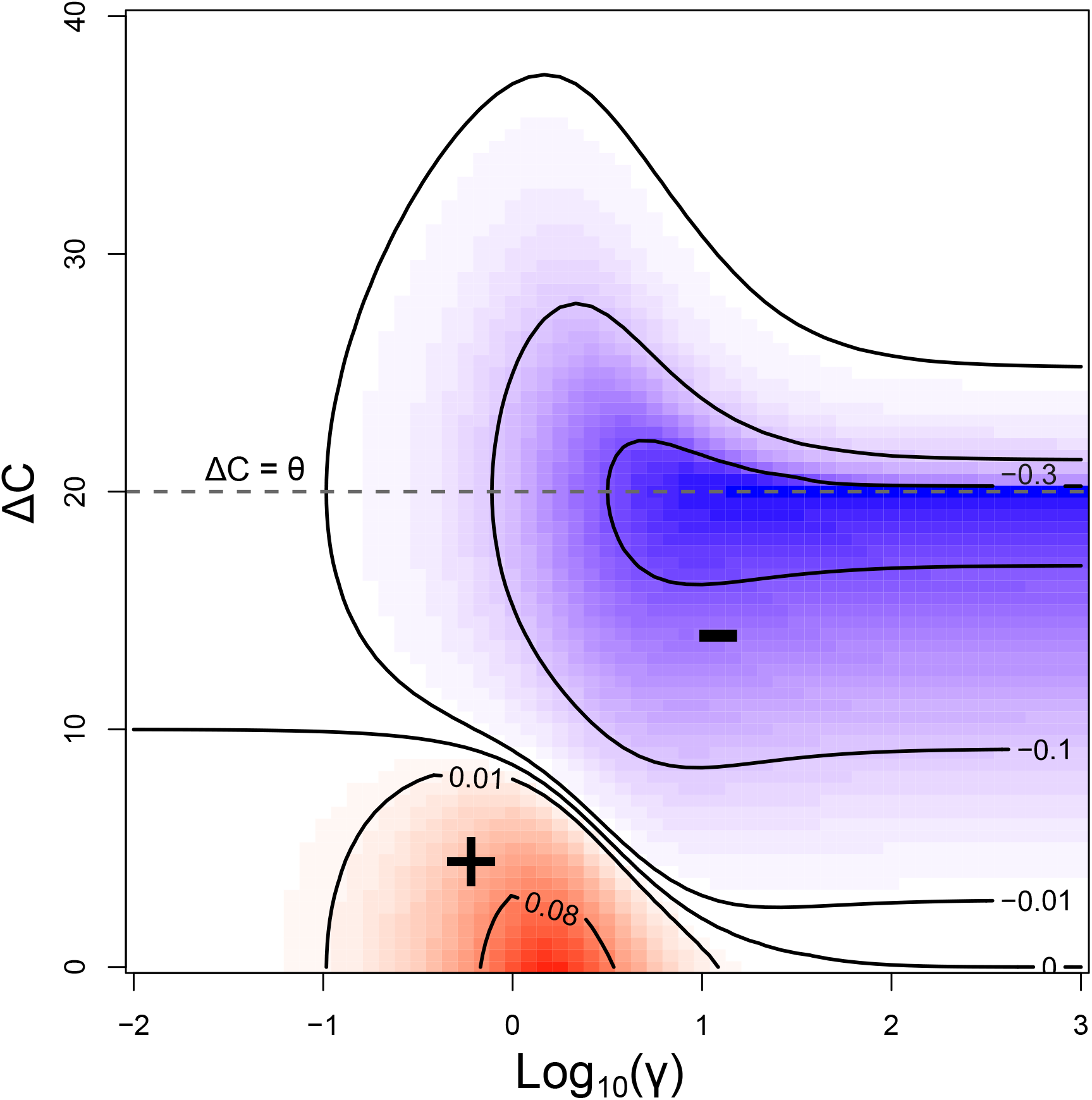
Emerging net species interactions as a function of the similarity of species’ environment optima (Δ*C*), and niche construction strength (*γ*). Parameter values for which net competitive interactions (−) are observed are depicted in blue, whereas net mutualistic interactions (+) are depicted in red. Interaction strength is measured by the relative change in the abundance of species 1 when associated with species 2, compared with its abundance when alone (Eq. 10). **Dashed line** Δ*C* = *θ* represents the threshold in environment optimum similarity that separates cases in which species’ contribution to the potential shows one or two minima. In the case where bistability is observed, we only study the equilibrium for which species 1 predominates (*C*_1_ = 40, *C*_2_ = *C*_1_ + Δ*C*, *E*_*t*=0_ = *B* = 50).

The diversity of ecosystem engineers also has an influence on system properties. In the case where species have weak niche construction abilities (*γ* = 1, Fig. 2 b), a single species is unable to create a well in the potential. Instead, the environment controls the dynamics and the only equilibrium corresponds to the environment baseline *B*. By contrast, when several weak engineer species with close optima are present, they are able to dig a common well in the potential landscape (Fig. 2, b, II). This leads to the emergence of an alternative stable equilibrium, in which the environment lies between the various species’ optima.

### 3.3 Influence of engineer similarity on species distribution and environmental changes in space

As described in section 3.2, the similarity of species environment optima (Δ*C*) influences the number of stable equilibria. When two ecosytem engineers are present along an environmental gradient, different community patterns can emerge, depending on Δ*C*. In the case where Δ*C > θ* (Fig. 4, I), each species pushes the environment to its own optimum. Along an environmental gradient, this leads to the emergence of distinct zones where the environment is driven close to the respective species optima. These zones are separated by abrupt changes in both the environment (Fig. 4,I,b) and species abundances (Fig. 4,I,c). Within these zones, each species is dominant in the spatial extent over which it controls the environment (Fig. 4,II). A distinct pattern emerges in the case where Δ*C < θ*, with the two species pushing the environment between their respective optima. This leads to the emergence of a single spatial zone where the environment is modified, and allows species coexistence at high abundances (Fig. 4, II, b-c). The transition between zones where the species can or cannot modify the environment is abrupt, with a discontinuity in both the environment and species abundances.

**Figure 4:**
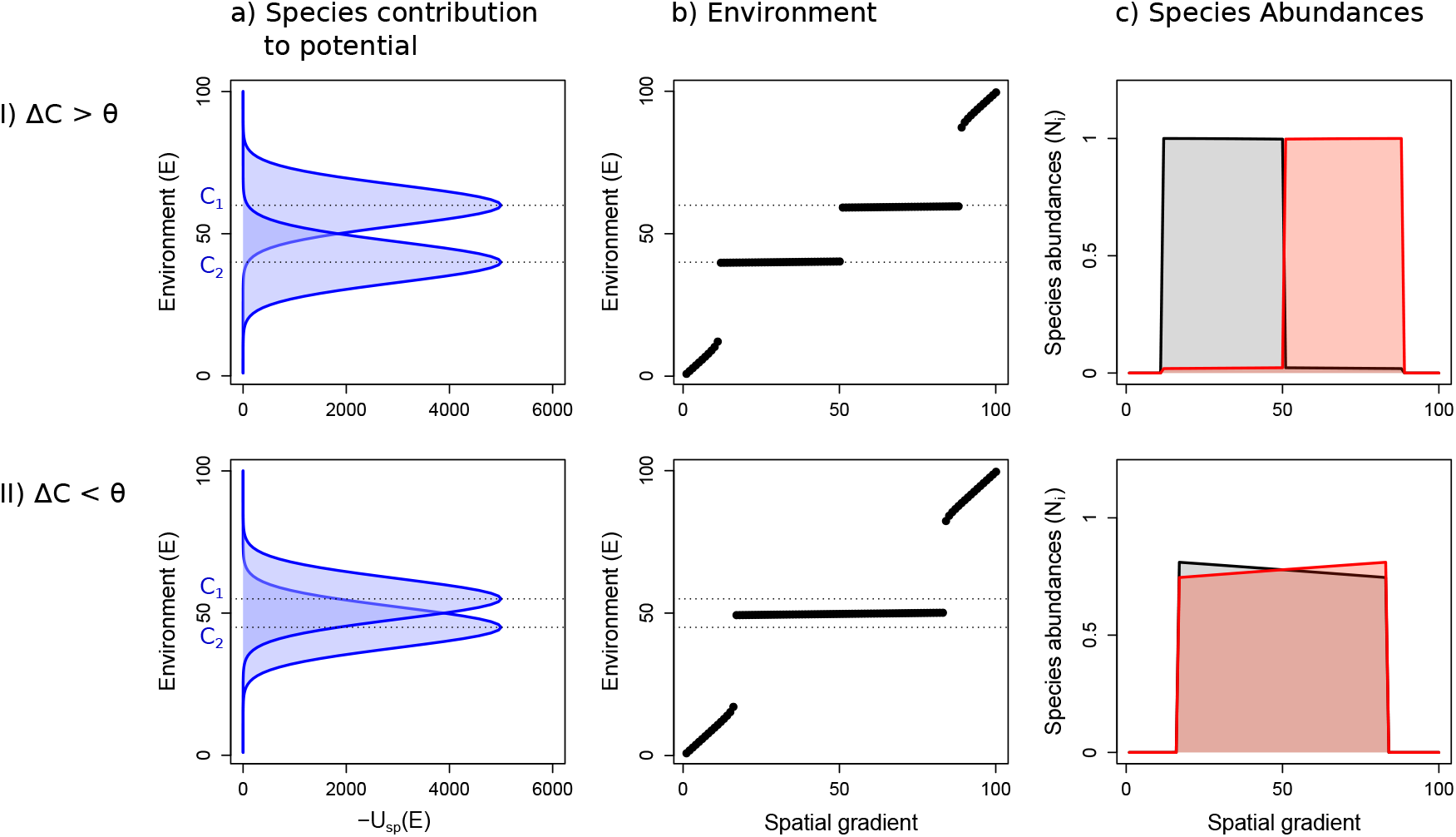
Influence of the similarity in ecosystem engineers on species distributions and the environment along a spatial gradient. We show results for: I) a difference in species’ environmental optima Δ*C* larger than the threshold *θ*, II) a difference in species’ environmental optima Δ*C* smaller than the threshold *θ*. (a): Species contribution - *U*_*sp*_(*E*) - to the potential *U* (*E*). (b): Value of the environment along the spatial gradient. (c): Species abundances along the spatial gradient at equilibrium. In the two depicted cases, species are strong ecosystem engineers (*γ* = 10).

### 3.4 Spatial community patterns in diverse communities

We now extend these results to many-species communities. In the case where several strong ecosystem engineers are present (*γ*_*i*_ = 10), we observe discrete communities in space, separated by sharp boundaries where important changes in both the abundance of ecosystem engineers (blue curves, Fig. 5, I) and in the environment (Fig. 5, I, b) occur. Non-engineers species (*γ*_*i*_ = 0, black curves) follow this pattern, with abrupt changes in their abundances. The bifurcation diagram shows the existence of alternative stable states, with different environment equilibria for a given location in space (Fig. 5, I, b). Similar patterns are observed when there are numerous weak ecosystem engineers (*γ* = 2), with the coincidence of abrupt changes in both the environment and species abundances in space. We observe much fewer discrete zones than there are engineers, because of the fusion of their potential wells (see section 3.2).

**Figure 5:**
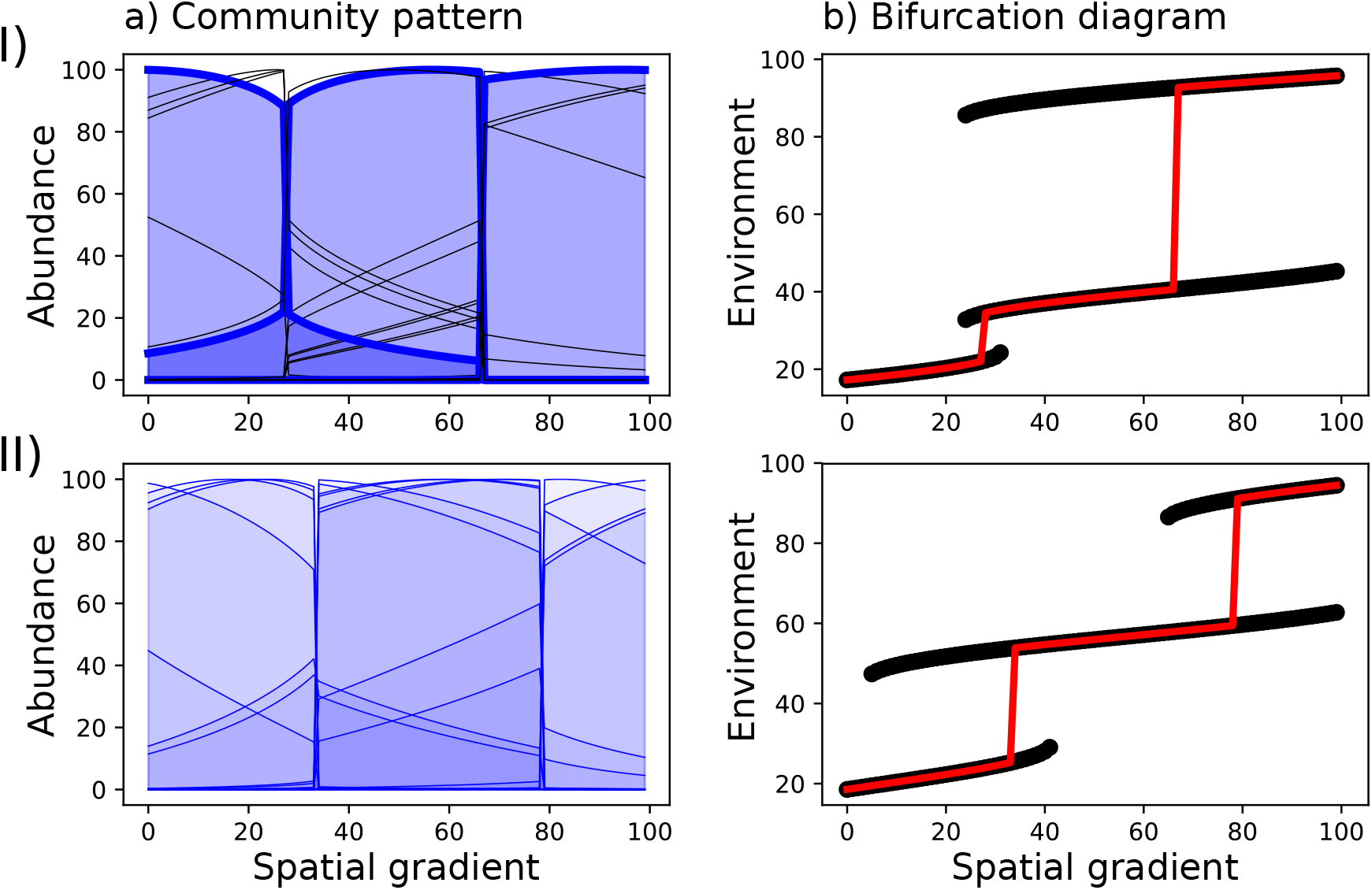
Species abundances along a spatial gradient (a) and bifurcation diagrams (b) in the case where: I) both strong ecosystem engineers (*γ* = 10, bold blue curves) and non-engineers (*γ* = 0, black curves) are present, II) numerous weak ecosystem engineers (*γ* = 2, blue curves) are present. In bifurcation diagrams (b), black curves represent all potential stable equilibria, and red lines represent equilibria observed in depicted cases in (a). Many weak engineers create fewer zones than there are engineers, and a pattern similar to the case where there are a few strong engineers.

## 4 Discussion

In this work, we investigated the role of biodiversity and species interactions in the emergence of ecotones through ecological niche construction. In particular, we studied the respective contributions of niche construction strength (*γ*), similarity in the environment optimum of the species (Δ*C*) and diversity (*S*). Our results show that, depending on the engineering strength *γ*, the contribution of biodiversity to ecotone emergence will be either through the similarity of species’ environmental optima Δ*C*, or through the diversity of engineering species *S*.

In the case of a single ecosystem engineer acting on the environment, discontinuities occur when a high niche construction rate (*γ*) allows the engineer to control its environment. These abrupt shifts are explained by the presence of two alternative stable states in the system that correspond to: 1) a modified state, with the environment close to the engineer’s optimum, and 2) a non-modified state, corresponding to the baseline value of the environment. A small change in the environmental conditions can thus lead to an abrupt shift from one attractor to the other.

In the case where species are strong ecosystem engineers, the difference in environmental optima (Δ*C*) is the main contribution of biodiversity to the emergence of ecotones. The presence of various engineers with distinct environment optima leads to the emergence of indirect interactions that influence the community patterns. We showed in a two-species system that these indirect interactions can be competitive or mutualistic, depending on the value of the difference Δ*C*.

When engineers have distant environmental optima and strong engineering abilities, their net interaction is competitive. At a given location, a species has a lower abundance when associated to a second engineer, as compared with its abundance when alone. Indirect competition through the environment can be observed in cases where there is multistability in the system, but also when a single equilibrium exists. In the extreme case where the modified environmental conditions are outside the other species’ fundamental niche, the latter can be excluded. By contrast, when the species’ environmental optima are close, with weak engineering abilities, we observe the emergence of net mutualistic interactions. In these cases, the two species are able to improve their carrying capacities, by modifying the environment to their mutual benefit. The abundance of a species is thus higher when associated with another engineer. In our study, the more species differ in their environmental optima, the stronger the negative effect they have on each other. This differs from classical limiting similarity theory (Abrams, 1983; MacArthur and Levins, 1967). Considering limiting resources such as water or light, limiting similarity theory predicts an increase in competition strength as the similarity in the resource requirements of the various species increases. By contrast, when species modify the abiotic environment to their own benefit, we showed that competition decreases, and then can turn into a net mutualistic interaction as the similarity of species’ environmental optima increases.

With more than two strong engineers along the gradient, engineers with close optima will tend to modify the environment to their collective benefit. When the ability of a community to modify the environment becomes higher than the ability of another one, the former will replace the latter along environmental gradients. This can be interpreted as a situation where there is competition between communities. In this case, the community shows a high level of integration (Clements, 1916; Wilson and Sober, 1989). This type of community organization tends to create particular species abundance patterns in space, with discrete communities separated by sharp boundaries.

In the case where the species are weak ecosystem engineers, the main contribution of biodiversity to community organization is through the number of engineering species. In this case, a weak ecosystem engineer alone is not able to substantially modify the environment and create a species-environment feedback. But when numerous weak engineers with similar optima are present, we do observe the emergence of species-environment feedbacks. In these cases, species jointly modify the environment to their collective benefit, as described above. In our model, an increase in species diversity can lead to an increase in each species’ biomass, through facilitation. The collective action of a large number of different ecosystem engineers can thus lead to the emergence of discrete communities along an environmental gradient, associated with sharp changes in the environment. In this study, the effect of several weak ecosystem engineers on the environment is not qualitatively different from the effect of a single strong engineer, but the spatial extent of the environmental change may be larger. The existence of several species may indeed broaden the spectrum of abiotic conditions under which the environment is modified, as seen in the case of positive interactions between two engineers. Biodiversity is potentially a key factor influencing the emergence of species-environment feedbacks in nature, and thus the emergence of sharp ecotones separating discrete communities. This might be the case in mangrove ecosystems, where several species can have similar effect on water salinity (Sternberg et al., 2007). As shown in this study, a certain level of biodiversity in ecosystem engineers might be necessary to maintain species-environment feedbacks. Likewise, Gonzalez et al. (2008) showed that the accumulation of small environmental changes by weak engineers can ultimately lead to a substantial change in the abiotic environment, and thus allow an ecosystem engineer to invade. A decrease in biodiversity, as currently observed worldwide (Pimm et al., 2014; Ceballos et al., 2015), might thus have important consequences, not only for community composition and organization, but also for the abiotic environment and for ecosystem functioning.

Species that do not modify their environment can also be influenced by ecological niche construction. By changing the environment, ecosystem engineers can promote species that benefit more from the modified state than the baseline conditions. In this case, ecosystem engineers indirectly facilitate other species through environmental modification. Facilitation has been shown to occur, particularly under harsh environmental conditions, such as in arid ecosystems (Soliveres and Maestre, 2014; Vega-Aélvarez et al., 2018; Armas and Pugnaire, 2005) or in cold environments (Choler et al., 2001; Callaway et al., 2002). When an engineer facilitates another species, it can be considered as a “nurse species” (Niering et al., 1963) that modifies the environment and allows the growth of species that would not have the ability to grow otherwise. Nevertheless, ecosystem engineering can also have negative effects on other species. For example, van Breemen (1995) showed how *Sphagnum* species can depress the growth of vascular plants by changing the environmental conditions in peat bogs ecosystems. A sharp ecotone can thus be explained by the appearance or disappearance of an engineer along the gradient, facilitating or preventing the growth of other species. In the case where species do not modify the environment to their own optimum, succession in time can be observed. In this case, the engineer can foster the growth of its successors, thus having a negative impact on its own performances (Appendix A4).

Species interactions - such as competition or mutualism - have been identified as drivers of species abundance along environmental gradients (Terborgh and Weske, 1975; Choler et al., 2001). We have shown in this paper that interactions between species and the abiotic environment can have unexpected consequences on species interactions themselves. These interactions can lead to the emergence of discontinuities in the environment, associated with sharp ecotones where important species turnover are observed. Explicit consideration of species-environment feedbacks is thus likely to increase our understanding of species distributions along environmental gradients. It may similarly be essential when studying the responses of species or communities to temporal changes in their environment. Finally, we have also shown that biodiversity can influence community organization along an environmental gradient. Current biodiversity loss can have major consequences for species distributions, abiotic environmental conditions, and ecosystem functioning.

## Data Accessibility Statement

Should the manuscript be accepted, the code supporting the results will be archived in an appropriate public repository and the data DOI will be included at the end of the article.

## Acknowledgements

We thank Yuval Zelnik for insightful discussions in preparing the manuscript, and two anonymous referees for their constructive contributions during revisions. This work was supported by the TULIP (Towards a unified theory of biotic interactions) Laboratory of Excellence (Project ANR-10-LABX-41), and by the BIOSTASES (BIOdiversity, STAbility and sustainability in Spatial Ecological and social-ecological Systems) Advanced Grant, funded by the European Research Council under the European Union's Horizon 2020 research and innovation program (grant agreement No 666971).

## Footnotes

1 But see Liautaud et al. (2019) for the role of dispersal in smoothing abrupt transitions.

