## Supporting Information for "Ecotone formation through ecological niche construction: the role of biodiversity and species interactions"

January 10, 2020

### A1 Overview

#### A1.1 Setting

When can we unambiguously identify the state of an ecosystem, demarcate its boundaries and follow its change over time? This is most easily done if its many component species, each with their individual goals and needs, behave and respond as a collective. Multiple mechanisms can plausibly lead to such large-scale self-organization in ecological communities.

A first possibility is a community of purpose, when all species have shared goals. This brings to mind the picture of a superorganism [1], a network of interactions carefully arranged toward self-perpetuation. But this picture has long been contested in ecology [2].

A second possibility is a community of means – a public good or marketplace tying together many individuals with different interests. When we can identify a large-scale ecosystem function, it is often tied to some shared resource – water, energy, basic elements. The existence of such a "common currency", through which a species can affect all others in a coherent fashion, is a widespread source of collective behavior.

This second possibility has already been largely explored in ecological theory, where resource competition plays a central role [3], and can indeed lead to collective organization [6]. Public goods models often assume that all agents benefit from accumulating some common resource; these works thus focus on setting up a tension between individual and collective means to achieve that profit.

Yet, the environment is no mere stockpile of resources: it is not only used, but also constructed and transformed. The environmental state – say, the concentration of various nutrients – differentially advantages some species over others, and the changes induced by a species need not be beneficial to itself. This can prompt many complex dynamics, such as a succession of different community stages, each benefitting from the outputs of the previous one.

#### A1.2 Model

We have $S$ species, each with abundance $N_i$, growth rate $r_i$ and interference competition $\alpha_{ij}$

$$\frac{dN_i}{dt} = r_i N_i \left(1 - \frac{N_i + \sum_j \alpha_{ij} N_j}{K_i(E)}\right) \tag{1}$$

Their carrying capacities are given by their environmental niche

$$K_i(E) = k_i^{\max} e^{-(E-C_i)^2/2T_i^2} \tag{2}$$

with a maximum of $k_i^{\max}$ when the environmental variable equals the species' optimum $E = C_i$, and a tolerance (niche width) of $T_i$. For numerical stability and clarity of interpretation, we can decide of an extinction threshold $\theta$ (e.g. a single individual) such that we treat smaller carrying capacities as being zero

$$K_i(E) < \theta \qquad \sim \qquad K_i(E) = 0. \tag{3}$$

Finally, each species affects the environment at a rate $m_i$ (the species' "engineering ability"), pushing it toward some value $\varepsilon_i$, while the environment tends to return to its baseline value $B$ with rate $\mu$,

$$\frac{dE}{dt} = \mu(B - E) + \sum_i m_i N_i (\varepsilon_i - E) \tag{4}$$

Perfect engineer species will have $\varepsilon_i = C_i$ and always draw the environment toward their own optimum. By contrast, imperfect engineers may create an environment which is suboptimal for themselves, $\varepsilon_i \neq C_i$, for instance by depleting resources that they need, or accumulating harmful byproducts.

To understand the long-term consequences of these dynamics, we will first study the equilibrium conditions

$$0 = N_i \left( K_i(E) - N_i - \sum_j \alpha_{ij} N_j \right)$$
$$E = \frac{\mu B + \sum_i m_i N_i \varepsilon_i}{\mu + \sum_i m_i N_i} \tag{5}$$

### A2 Perfect coexisting engineers

Throughout this section, we assume perfect engineer species ($\varepsilon_i = C_i$) without interference competition, $\alpha_{ij} = 0$. Then, all species (with nonzero carrying capacity) can coexist, and at equilibrium

$$N_i = K_i(E). \tag{6}$$

#### A2.1 Potential landscape and equilibria

##### A2.1.1 Slow environment

If the dynamics of species abundances is much faster than that of the environment $r_i \gg m_i, \mu$, we expect that species quickly reach their carrying capacity for a given environment value, $N_i = K_i(E)$, and hence the dynamics of the system is given by

$$\frac{dE}{dt} = \mu(B - E) + \sum_i m_i K_i(E)(C_i - E) \tag{7}$$

We now show that is in fact a *gradient descent* dynamics, similar to

$$\frac{dx}{dt} = -\frac{dU(x)}{dx} \tag{8}$$

where $U(x)$ is a potential function, with the dynamics always going toward the closest minimum of $U(x)$.

Indeed, notice that

$$K_i(E)(C_i - E) = T_i^2 \frac{d}{dE} K_i(E) \tag{9}$$

Thus,

$$\frac{dE}{dt} = -\frac{d}{dE} U(E) \tag{10}$$

where the potential takes the form:

$$U(E) = \frac{\mu}{2}(E - B)^2 - \sum_i m_i T_i^2 K_i(E) \tag{11}$$
$$= \frac{\mu}{2}(E - B)^2 - \sum_i m_i k_i^{\max} T_i^2 e^{-(E-C_i)^2/2T_i^2} \tag{12}$$

We see it has two components: a parabolic well $(E - B)^2$ which has a single minimum at $B$, and a sum of Gaussian wells created by each of the engineer species. If $m_i$ are large enough, $U(E)$ can have local minima corresponding to these engineered wells, and if $m_i \gg \mu$, these wells are deeper than the parabola, so engineered states are more stable than the natural state.

The effective strength of a species' long term action on the environment is thus

$$\lambda_i = m_i k_i^{\max} T_i^2 \tag{13}$$

meaning that a species can be an important ecosystem engineer either through large engineering ability $m_i$, large maximum abundance $k_i^{\max}$, or wide niche $T_i$.

At a given patch, as $E(t)$ changes over time, $U(E(t))$ will decrease until it reaches the bottom of the local basin:

$$\frac{dU}{dt} = \frac{dU}{dE}\frac{dE}{dt} = -\left(\frac{dE}{dt}\right)^2 \leq 0 \tag{14}$$

See Fig. 4 for a map of the potential in a simulation and how it can be used to predict the final environmental state.

###### A2.1.2 Fast environment

We now consider the opposite limit, when the dynamics of the environment variable $E$ are faster than the species'. Crucially, equilibria are independent from the relative timescale of environment and species dynamics. Therefore, equilibria must always be minima of the potential $U(E)$, even for fast environment dynamics. However, in that case, there is no guarantee that these dynamics can be approximated by gradient descent, meaning that $E(t)$ will not necessarily remain within the initial basin of attraction.

To move out of the initial basin, there must be some time during which the dynamics are climbing up the potential landscape, i.e. $dU/dt > 0$. Notice that

$$\frac{dU}{dt} = \frac{dU}{dE}\frac{dE}{dt} = \frac{dU}{dE}\left(-\frac{dU}{dE} + \sum_i m_i(N_i - K_i(E))(C_i - E)\right) \tag{15}$$

hence $dU/dt > 0$ requires the second term in parentheses to be larger than the slope $dU/dE$ of the potential. Given a potential barrier, the dynamics may climb it if:

- there exist basins beyond the barrier (in the same direction as the slope, i.e. same sign of $dU/dE$ and $C_i - E$) that are created by species with $N_i > K_i(E)$
- there exist basins in the other direction (opposite sign of $dU/dE$ and $C_i - E$) created by species with $N_i < K_i(E)$

In other words, faster environmental dynamics will be able to climb out of shallow wells provided that there are overabundant species attracting them, or, more likely for barren initial conditions, if the species creating these wells remain at low abundance $N_i < K_i(E)$ for sufficiently long. Thus, the dynamics are most likely to settle in a deep (and not necessarily wide) basin. By contrast, slow environment dynamics might favor the widest basin, which is more likely to contain the initial condition $E(0)$ (see Discussion).

###### A2.1.3 Effective species interactions for fast environment

Let us set $\mu = 1$ for simplicity (i.e. $m_i$ is measured in units of $\mu$). Then the equilibrium environment value is

$$E = \frac{B + \sum_i m_i N_i C_i}{1 + \sum_i m_i N_i}$$

If the environment quickly reaches this value for any species abundance $N_i$, the dynamical equation for species becomes

$$\frac{1}{r_i N_i}\frac{dN_i}{dt} = 1 - \frac{N_i}{k_i^{\max}} \exp\left[\frac{1}{2T_i^2}\left(\frac{(B - C_i) + \sum_j m_j N_j (E_j - C_i)}{1 + \sum_j m_j N_j}\right)^2\right] \tag{16}$$

For $m_i \ll 1$, we can do a series expansion

$$\frac{(B - C_i) + \sum_j m_j N_j (E_j - C_i)}{1 + \sum_j m_j N_j} \approx (B - C_i) + \sum_j m_j N_j (E_j - B) \tag{17}$$

and thus

$$\frac{1}{r_i N_i} \frac{dN_i}{dt} \approx 1 - \frac{N_i}{K_i(B)} \left( 1 + \frac{1}{T_i^2} \sum_j m_j N_j (B - C_i)(E_j - B) \right). \tag{18}$$

At equilibrium

$$N_i = \frac{K_i(B)}{1 + \frac{1}{T_i^2} \sum_j m_j N_j (B - C_i)(E_j - B)} \tag{19}$$

Once again, we do a Taylor expansion to get

$$N_i = K_i(B) \left( 1 - \frac{1}{T_i^2} \sum_j m_j N_j (B - C_i)(E_j - B) \right) \tag{20}$$

which is equivalent to the equilibirum of a Lotka-Volterra model

$$N_i = K_i(B) \left( 1 - \sum_j A_{ij} N_j \right), \qquad A_{ij} = \frac{d \log K_i}{dN_j}(B) = \frac{m_j}{T_i^2}(B - C_i)(E_j - B). \tag{21}$$

#### A2.2 Effective facilitation and competition

##### A2.2.1 When does an engineer species create an equilibrium?

Let us consider a single engineer species and ask when it can create an equilibrium with a value of $E$ distinct from $B$. The equilibrium criterion is $dU/dE = 0$, i.e.

$$\mu(E - B) = m_1 k_1 (C_1 - E) e^{-(E - C_1)^2 / 2T_1^2} \tag{22}$$

For simplicity, let us consider the case $C_1 - B > T_1$. Then, we can approximate our question by asking whether the maximum of the right-hand term at some particular value $E_m$ is larger in absolute value than the left-hand term $\mu(E_m - B)$ at that point. The maximum of the right-hand term is given by a zero of its derivative

$$m_1 k_1 \left( \frac{(E_m - C_1)^2}{T_1^2} - 1 \right) e^{-(E_m - C_1)^2 / 2T_1^2} = 0 \tag{23}$$

hence $E_m = C_1 - T_1$, and a sufficient condition for species 1 to create a new equilibrium is

$$\mu(C_1 - B - T_1) \lesssim m_1 k_1 T_1 e^{-1/2}. \tag{24}$$

We see that it is harder to create a new equilibrium far away from the baseline environment value, i.e. when $C_1 - B$ is large.

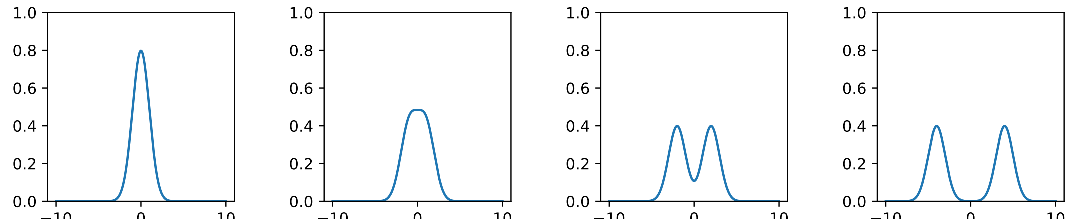

Figure 1: Joint effect of two species on the potential landscape: sum of two Gaussian functions of equal standard deviation $T$, separated by $\Delta C \in \{0, 2, 4, 8\}$ (left to right). Up to $\Delta C = 2T$, the sum of Gaussians is unimodal, indicating that the two species can create a single equilibrium together. Afterward, species are effectively in competition, as each one prevails in a different equilibrium state. Still, until $\Delta C \approx 8T$, the overlap increases the height of each peak, while beyond, there is little to no positive effect of one species on the other.

##### A2.2.2 Two species

To understand the long-term interactions between two species through their engineering capabilities, we can study the environmental variable's potential landscape $U(E)$ and ask: does each species create a potential well (are there as many alternative stable states as there are species)?

The contribution of engineers to $U(E)$ is a sum of Gaussian terms,

$$U(E) = \frac{\mu}{2}(E - B)^2 - \sum_i \lambda_i e^{-(E-C_i)^2/2T_i^2} \tag{25}$$

where $\lambda_i$ is defined in (13). If we assume for now $\mu = 0$, our question becomes: when is the sum of two Gaussians unimodal?

Consider two species that are equal in every respect, save their optimum

$$\lambda_1 = \lambda_2, \quad T_1 = T_2 = T, \quad C_2 - C_1 = \Delta C \tag{26}$$

We find three cases (see Fig. 1)

- Pure competition: if $\Delta C \gg 2T$, the two Gaussians are well separated and each species forbids the other from existing
- Mixed facilitation and competition: if $\Delta C \gtrsim 2T$ (e.g. $\Delta \in [2T, 8T]$), the two species allow each other to exist, and even facilitate each other to some extent (making the other's potential well deeper, and thus more likely to overcome environment inertia or other competitors). Still, the sum of Gaussians remains bimodal: there are two possible equilibria, each favoring one of the species. This is a form of moderate competition between facilitators.
- Coalescence: if $\Delta C < 2T$, the sum of Gaussians becomes unimodal, with a peak halfway between the species optima. Now, the two species act together as a single, more influential species.

If species differ in other parameters, the same qualitative picture holds, although the weaker species (smaller $\lambda$) will need a larger $\Delta C$ to maintain its own distinct peak, rather than be absorbed in the stronger species'. There exists a general quantitative criterion for bimodality in a mixture of two arbitrarily different Gaussians [5], which we report in the next section, but it does not easily generalize to more species.

##### A2.2.3 Criterion for bimodality of a Gaussian mixture

According to [5], a weighted sum of two gaussians

$$pG(x, 0, 1) + (1-p)G(x, \mu, \sigma) \tag{27}$$

is bimodal if $\mu > \mu_0$ with

$$\mu_0 = \frac{1}{\sigma}\sqrt{2(\sigma^4 - \sigma^2 + 1)^{3/2} - (2\sigma^6 - 3\sigma^4 - 3\sigma^2 + 2)} \tag{28}$$

and $p \in [p_1, p_2]$ where

$$\frac{1}{p_i} = 1 + \frac{\sigma^3 y_i}{\mu - y_i} e^{-\frac{1}{2}y_i^2 + (y_i - \mu)^2/(2\sigma^2)} \tag{29}$$

with $y_1$ and $y_2$ the roots of the equation

$$(\sigma^2 - 1)y^3 - \mu(\sigma^2 - 2)y^2 - \mu^2 y - \mu\sigma^2 = 0 \tag{30}$$

with $0 < y_1 < y_2 < \mu$. Otherwise, the sum is unimodal.

##### A2.2.4 Many species

For many species, no exact results exist but we can provide a scaling estimate of the average number of alternative states. Given $S$ the number of species, with their optima distributed over interval $[0, L]$, the average distance between their optima is

$$\langle \Delta C \rangle = L/S. \tag{31}$$

If the optima are uniformly distributed, the number of optima within a certain interval follows a Poisson distribution, and thus, the probability that $\Delta C > 2T$ (avoiding coalescence) is the probability of having no optima within a span of $2T$, i.e.

$$P(\Delta C > 2T) \approx e^{-2\langle T \rangle / \langle \Delta C \rangle} \tag{32}$$

Hence, the typical number of clusters of coalesced species scales like

$$S_0 \sim S e^{-2\langle T \rangle S/L}. \tag{33}$$

This approximation will only hold up to $\langle \Delta C \rangle \sim T$, i.e. $S \sim L/2\langle T \rangle$, after which adding more species will typically not contribute more equlbria.

Now recall that species $i$ on its own can create an alternative equilibrium despite the natural environmental dynamics only if

$$\mu(C_i - B) \lesssim m_i k_i^{\max} T_i \tag{34}$$

meaning that the potential well created by the species is deep enough to compensate the recovery of the environment, which gets faster as $E$ moves away from $B$. This means that species can only contribute to an equilibrium if their optimum falls within a range

$$L' \leq \min\left(L, \frac{2\langle mkT \rangle}{\mu}\right). \tag{35}$$

Only the fraction $L'/L$ of species clusters with optima within that range can create new equilibria.

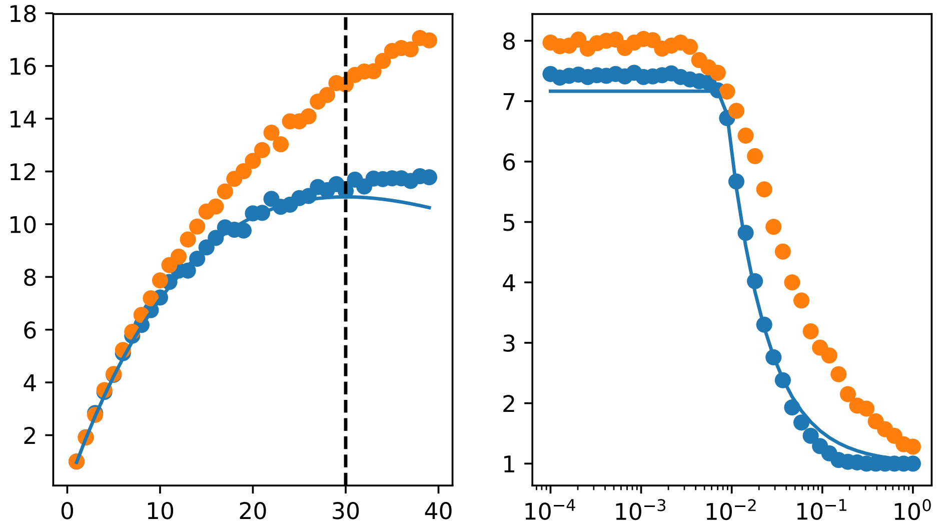

Figure 2: Number of alternate stable states $N_{eq}$ as a function of number of species $S$ (left panel) and environment recovery $\mu$ (right panel). Species have identical niches (blue dots) or heterogeneous niche widths $T_i$ drawn from a Gamma distribution with variance 0.3 (orange dots). The solid lines represent the analytical prediction (36). The dashed line indicates $S = L/2\langle T\rangle$, the threshold above which $N_{eq}$ saturates.

Therefore, the expected number of equilibria (including the natural equilibrium at $B$) is

$$\begin{aligned} N_{eq} &\sim 1 + S_0 \frac{L'}{L} \\ &\lesssim 1 + S e^{-2\langle T\rangle S/L} \min\left(1, \frac{\langle mkT\rangle}{\mu L}\right). \end{aligned} \tag{36}$$

This simple formula reproduces the qualitative behaviors within a wide range of parameters, see Fig. 2. Deviations happen if species are heterogeneous in their properties $k_i^{\max}$, $m_i$ or $T_i$, and as mentioned above, our calculation does not account for the saturation beyond a threshold $S > L/2\langle T\rangle$ shown by the dashed line in Fig. 2.

#### A3 Interference competition and imperfect engineers

Let us now consider the case of direct interference competition $\alpha_{ij} \neq 0$ and imperfect engineers $\varepsilon_i \neq C_i$. Now, some species can go extinct even while their carrying capacity is nonzero. Recall the equilibrium condition

$$0 = N_i \left( K_i(E) - N_i - \sum_j \alpha_{ij} N_j \right)$$

The abundance of surviving species $N_i^* \neq 0$ is given by

$$N_i^* + \sum_j \alpha_{ij} N_j^* = K_i(E) \tag{37}$$

or in vector form,

$$N^* = (\mathbb{I} + \alpha^*)^{-1} K^*(E) \tag{38}$$

where $\mathbb{I}$ is the identity matrix, $\alpha^*$ is the matrix of interactions restricted to the $S^*$ surviving species, and $K^*(E)$ the vector of carrying capacities of surviving species. For convenience, define the matrix

$$V_{ij} = (\mathbb{I} + \alpha^*)^{-1}_{ij} \tag{39}$$

so that

$$N^* = V K^*(E). \tag{40}$$

#### A3.1 Equivalence

Taking once again the limit of fast species dynamics, we now have

$$\begin{aligned}\frac{dE}{dt} &= \mu(B - E) + \sum_{ij} m_i(\varepsilon_i - E) N_i^* \\ &= \mu(B - E) + \sum_{ij} m_i(\varepsilon_i - E) V_{ij} K_j(E)\end{aligned} \tag{41}$$

which we can rewrite as

$$\frac{dE}{dt} = \mu(B - E) + \sum_i^{S^*} \hat{m}_i(\hat{\varepsilon}_i - E) K_i(E) \tag{42}$$

with

$$\hat{m}_i = \sum_j^{S^*} V_{ji} m_j, \qquad \hat{\varepsilon}_i = \frac{\sum_j^{S^*} V_{ji} m_j \varepsilon_j}{\sum_j^{S^*} V_{ji} m_j} \tag{43}$$

Thus, we see that direct competition appears equivalent, in terms of its equilibrium effect, to imperfect engineering with effective values of species engineering capability $m_i$ and target environment value $\varepsilon_i$. An important consequence is that even perfect engineers ($\varepsilon_i = C_i$) will behave like imperfect ones if they also interact directly.

It is, however, crucial to note that the calculation above involves summing only on the $S^*$ species that survive the direct competitive interaction. While the matrix $V_{ij}$ depends only on interactions $\alpha_{ij}$ and not on the environment, it does depend on who survives, which is controlled by the carrying capacities as well. Thus, direct competition cannot simply be replaced by imperfect engineering, except in the regime where all species coexist (i.e. for weak direct interactions). In particular, if direct competition allows for alternate equilibria (mutual exclusion), each will correspond to a different equation (42).

Perhaps counter-intuitively, it is the action of species $i$ on others that appears in the effective parameters above. For instance, if direct interactions are weak,

$$V = (\mathbb{I} + \alpha^*)^{-1} \approx \mathbb{I} - \alpha^* \tag{44}$$

$$\hat{m}_i \approx m_i - \sum_j^{S^*} \alpha_{ji} m_j, \qquad \hat{\varepsilon}_i \approx \varepsilon_i - \sum_j^{S^*} m_j \alpha_{ji}(\varepsilon_j - \varepsilon_i) \tag{45}$$

and we see that a species' effective engineering ability $\hat{m}_i$ decreases due to its competitive effect on others, $\alpha_{ji}$, while its effective target environmental value $\hat{\varepsilon}_i$ moves away from the optima $\varepsilon_j$ of the species it affects.

#### A3.2 Skewed potential

From equation (42), we can construct the corresponding potential by noticing that

$$\frac{dE}{dt} = -\frac{dU(E)}{dE} = \mu(B-E) + \sum_i^{S^*} \hat{m}_i (C_i - E) K_i(E) + \sum_i^{S^*} \hat{m}_i (\hat{\varepsilon}_i - C_i) K_i(E) \tag{46}$$

The last term is the only one that differs significantly from the equation in the case of perfect engineers, (7). Thus, we can separate the resulting potential into two contributions: first, the usual potential for perfect engineers, obtained here with the $S^*$ surviving species and effective engineering rates $\hat{m}_i$, and second, a correction $\Delta U(E)$ coming from the last term above. We write

$$U(E) = U_{pe}(E) + \Delta U(E) \tag{47}$$

where

$$U_{pe}(E) = \frac{\mu}{2}(E-B)^2 - \sum_i^{S^*} \hat{m}_i T_i^2 K_i(E) \tag{48}$$

is the contribution that is similar to the perfect engineer case. Since $K_i(E)$ is Gaussian, its integral is an error function, and the correction to the potential takes the form

$$\Delta U(E) = \sum_i^{S^*} \hat{m}_i T_i^2 (\hat{\varepsilon}_i - C_i) \operatorname{erf}\left( \frac{E - C_i}{\sqrt{2} T_i} \right) \tag{49}$$

Error functions are sigmoidal and comprised between 0 and 1, so $\Delta U(E)$ will have the general shape of a "staircase", i.e. a sum of step-like functions going up or down, with a step height of $\hat{m}_i T_i^2 (\hat{\varepsilon}_i - C_i)$.

This could create new potential wells, if two (or more) species push $E$ in the direction of each other's optimum, giving rise to a new type of interaction: obligate facilitation (or mutual stabilization), where each species degrades its environment from its own perspective, but improves it from the perspective of the other.

If $\hat{\varepsilon}_i - C_i$ has the same sign for many species (e.g. all species tend to degrade complex sugars into simpler ones, pushing the environment variable $E$ in a constant direction), the effect will be to create a general slope in that direction, and thus, dynamics akin to succession.

### A4 Ecotones and succession

#### A4.1 Ecotones on an environmental gradient

Let us assume an environmental (e.g. latitudinal or altitudinal) gradient, reflected in the fact that the baseline value of the environmental variable $B = B(x)$ now depends on position $x$ along the gradient.

If we choose a solution of (5) and follow it along the gradient, as we progressively change parameters such as $B(x)$, we do not expect sharp ecotones (transition zones between communities with different species compositions and abundances). The only option for a singular transition is to have alternative stable states, with the transition occurring when one of the states loses its stability.

Whenever two or more attractors exist for the same patch $x$, see black lines on Fig. 4 top-right, there is potential for hysteresis.

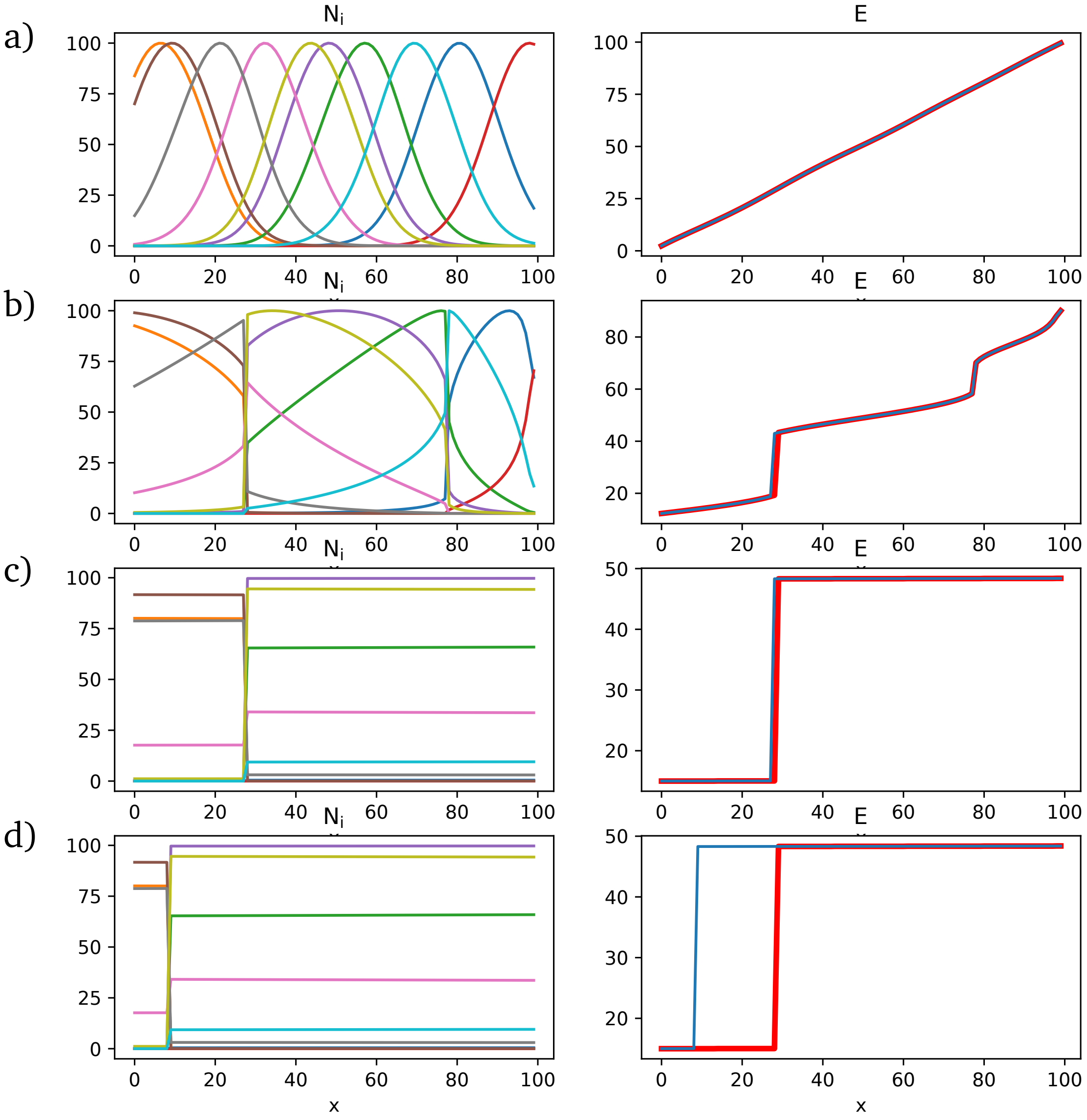

Figure 3: Realized abundance for species in the community (left) and values of the environmental variable (right) along a spatial gradient defining the baseline environmental variable $B(x) = x$ which is also the initial state at each point. Increasing $\bar{m}/\mu$ from 0.1 (a) to 10 (b) to 100 (c), we go from an environment that closely follows $B$, and thus a continuous turnover of species, to the existence of alternate stable states engineered by these species, and separated by sharp transitions. Finally, reducing the rate of species dynamics from $r = 10$ (c) to $r = 0.1$ (d) increases the difference between the pure gradient descent prediction (red line, right panels) and the observed environment value $E$ at each position $x$. Possible equilibria are independent of $r$, but which equilibrium is reached does depend on it: as we explain in Fig. 4, high $r$ entails gradient descent toward the closest equilibrium, while low $r$ allows the dynamics to climb a potential wall, and tends to favor the deeper basins rather than the ones closest to the initial condition.

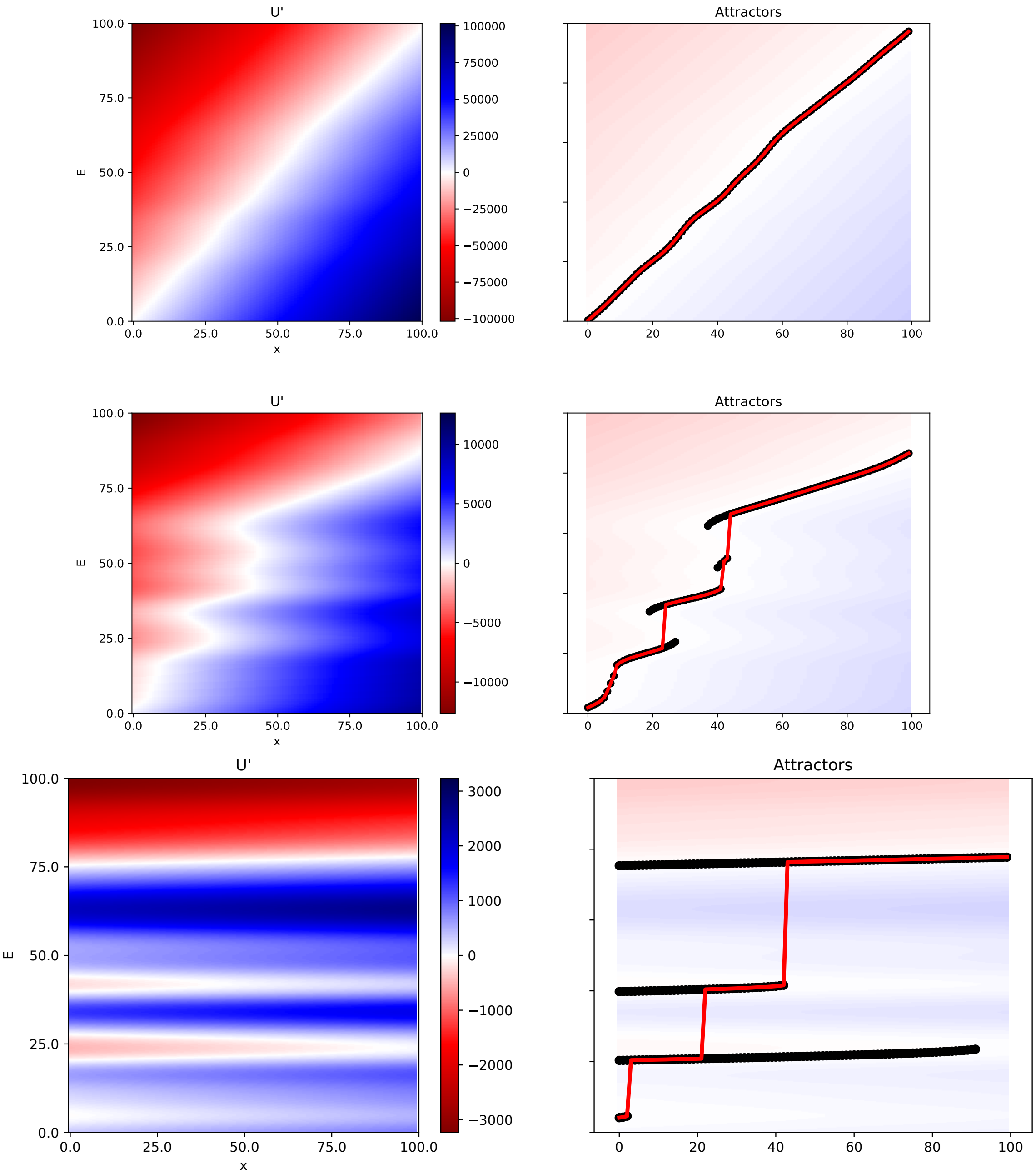

Figure 4: Using the potential landscape $U(E)$ to predict the equilibrium state, for $\mu = 1000, 100, 1$ (top to bottom). Left column: $dU/dE$ as a function of the environment variable $E$ (y-axis) and the position $x$ on the gradient (x-axis) which controls the natural environment value $B(x) = x$. Equilibria correspond to white lines $dU/dE = 0$ (stable if red is above and blue below, or unstable the other way around). Right column. Following the stable equilibria (black dots) and predicting where gradient descent should go if $E(x, t = 0) = B(x)$ (red line).

##### A4.2 Succession trajectories

In this model, including direct competition and imperfect engineering, one can imagine three different succession scenarios.

The reference scenario is the usual competition-colonization tradeoff setting [7]. In that case, species at later stages are expected to have slower growth but stronger competitive ability, either through direct competition, engineering ability, or a combination of both. As a consequence, later stages will be longer, but transitions between stages will also be slower.

Another scenario is succession driven by the environmental variable slowly descending down the potential landscape created by the engineer species. In that case, there is no implication that later stages will be longer. If the landscape is shaped by perfect engineers, we can expect a rather smooth change of the environmental variable. If it is shaped by imperfect engineers, it is possible to have long transition periods where all species have low abundance, separating shorter periods where a set of imperfect engineers dominates. There may be stabilization at low abundances if two sets of engineers are pushing the environment toward each other.

The third scenario is perturbation-driven succession: jumps between equilibria, either due to random noise, or to directed perturbations (e.g. a gradual increase in the baseline environmental value $B$). It is only in this scenario that succession will generically exhibit discrete stages separated by sudden transitions. Under random perturbations, succession will proceed on average toward deeper wells, and thus later stages will be longer on average (deeper wells resist perturbations longer), but there may be reversions to earlier stages.

#### A5 Supplementary Discussion

##### A5.1 Slow and fast environment

An intuitive aspect of the speed of environment change is its inertia upon removal of some engineer species. In one limit, the environmental state may remain the same for long times (for instance, peat created by Sphagnum mosses can remain for thousands of years [8]), long enough that the species could potentially recolonize at the same abundance before any significant change occurred. In the other limit, the environmental state may revert suddenly, even instantaneously when the engineering results from physical properties of the species themselves (e.g. shielding of light by the canopy).

As noted in Sec. A2, when the environment dynamics are slow, the environmental variable effectively follows a gradient descent. This means that the species (or group of species) creating the widest basin of attraction control the dynamics for a broad range of initial conditions. By contrast, when the environment dynamics are fast, they are drawn toward the optimum of the species with the largest carrying capacity and best engineering abilities, even if its niche is narrow. We thus predict a prevalence of generalists in slow environments, and specialists in fast environments.

##### A5.2 Multiple environmental variables

A single environmental variable $E$ may not suffice to accurately represent the ways in which species interact through modifications of their surroundings. On the other hand, our modelling approach for ecosystem engineers is most relevant if the number of environmental variables is limited, and small compared to the number of species – otherwise, it may be simpler to directly model pairwise species interactions.

There is one important qualitative feature that distinguishes the outcomes of this model, and those of a model with multiple environmental "dimensions": with only one dimension $E$, there can be at most as many equilibria as there are engineer species. This stops being the case with more environmental variables. A corresponding mathematical result states that a mixture of Gaussian components can have more maxima than components in dimension $d > 1$ [4, 9]. This means that some equilibria could not be assigned to, nor expected from, the action of any given species on its own.

[9] Benjamin Wallace. *On the critical points of Gaussian mixtures*. PhD thesis, Queen's University, Kingston, Ontario, Canada, 2013.
